# Structural basis for SARS-CoV-2 Nucleocapsid protein recognition by single-domain antibodies

**DOI:** 10.1101/2021.06.01.446591

**Authors:** Qiaozhen Ye, Shan Lu, Kevin D. Corbett

## Abstract

The COVID-19 pandemic, caused by the coronavirus SARS-CoV-2, is the most severe public health event of the twenty-first century. While effective vaccines against SARS-CoV-2 have been developed, there remains an urgent need for diagnostics to quickly and accurately detect infections. Antigen tests, particularly those that detect the abundant SARS-CoV-2 Nucleocapsid protein, are a proven method for detecting active SARS-CoV-2 infections. Here we report high-resolution crystal structures of three llama-derived single-domain antibodies that bind the SARS-CoV-2 Nucleocapsid protein with high affinity. Each antibody recognizes a specific folded domain of the protein, with two antibodies recognizing the N-terminal RNA binding domain and one recognizing the C-terminal dimerization domain. The two antibodies that recognize the RNA binding domain affect both RNA binding affinity and RNA-mediated phase separation of the Nucleocapsid protein. All three antibodies recognize highly-conserved surfaces on the Nucleocapsid protein, suggesting that they could be used to develop affordable diagnostic tests to detect all circulating SARS-CoV-2 variants.

## Introduction

The COVID-19 pandemic, caused by the novel coronavirus SARS-CoV-2, has resulted in over 170 million confirmed cases and caused over 3.5 million deaths as of early June 2021 (John Hopkins Coronavirus Resource Center, https://coronavirus.jhu.edu). Despite the development and widespread use of effective vaccines against SARS-CoV-2 (Polack et al., 2020; Baden et al., 2021; Sadoff et al., 2021), the virus and its emerging variants will remain an active public health threat for the foreseeable future. In order to detect and quickly respond to new infections and local outbreaks, robust diagnostic tools are required that can quickly and reliably detect active SARS-CoV-2 infections.

Antigen tests are immunoassays that detect the presence of a viral antigen such as an abundant protein, and constitute an effective means of detecting active infections for respiratory pathogens including SARS and SARS-CoV-2 (Che et al., 2004; Guglielmi, 2020; Sethuraman et al., 2020). Antigen tests are an important complement to PCR-based tests, which detect the presence of viral nucleic acids (genomic or sub-genomic RNA in the case of SARS-CoV-2), but may give positive results after a patient is no longer infectious. While antigen tests are less sensitive than PCR-based tests, they are generally faster and require less specialized equipment than PCR tests, enabling wider deployment than PCR-based tests.

The major structural proteins of SARS-CoV-2 include S (Spike), M (Membrane), E (Envelope), and N (Nucleocapsid). While the Spike protein is exposed on the surface of virions and is the target of all major vaccines, the Nucleocapsid protein is highly abundant in virions and infected cells, and is therefore a common choice for antigen tests. The N protein plays several roles in the SARS-CoV-2 life cycle, including facilitating viral RNA production, suppressing host cells’ innate immune responses, and packaging viral genomic RNA into developing virions. To accomplish these tasks, the N protein possesses a modular structure with an N-terminal RNA-binding domain (RBD) and a C-terminal dimerization domain (CTD), plus three intrinsically disordered regions (IDRs) at the N- and C-termini and between the RBD and CTD. The protein oligomerizes through its CTD and disordered C-terminal tail (Ye et al., 2020), and the protein also undergoes liquidliquid phase separation with RNA mediated by its RBD and central disordered region (Carlson et al., 2020; Cubuk et al., 2020; Iserman et al., 2020; Perdikari et al., 2020; Savastano et al., 2020; Lu et al., 2021; Luo et al., 2021). In cells, N protein condensates recruit the stress granule proteins G3BP1 and G3BP2, suppressing stress granule assembly (Nabeel-Shah et al., 2020; Lu et al., 2021; Luo et al., 2021). During virion production, the N protein assembles into viral RNA-protein complexes (RNPs) with a characteristic barrel shape to package the viral RNA (Klein et al., 2020; Yao et al., 2020), and interacts with the Membrane protein to recruit the viral genome to developing virions (Lu et al., 2021).

Two recent studies reported the isolation of a total of four single-domain antibodies (sdAbs) that target SARS-CoV-2 N with high affinity (Anderson et al., 2021; Sherwood and Hayhurst, 2021). Here, we map the epitopes of these sdAbs and show that each sdAb recognizes a specific domain of SARS-CoV-2 N. We report high-resolution crystal structures of two sdAbs bound to the N RBD, and of one sdAb bound to the N CTD. Comparison of the mapped sdAb epitopes with N protein mutations in a set of nearly 500,000 SARS-CoV-2 genomes shows that the sdAbs recognize surfaces that are highly conserved across SARS-CoV-2 isolates and in major variants of concern, suggesting broad utility in recognizing SARS-CoV-2 infections.

## Results

### sdAbs recognize distinct domains of the SARS-CoV-2 N protein

Two recent studies reported the isolation of single-domain antibodies (sdAbs) that recognize the SARS-CoV-2 N protein (**Figure 1a-b**) (Anderson et al., 2021; Sherwood and Hayhurst, 2021). One study used an in vitro selection method to isolate one sdAb (here termed sdAb-N3; **Table S1**) that recognizes the N protein with high affinity (EC_50_ ~50 nM as measured by luciferase-based ELISA assay) (Sherwood and Hayhurst, 2021). The second study isolated three sdAbs (here termed sdAb-B6, sdAb-C2, and sdAb-E2; **Table S1**) from an immunized llama, which each recognize the N protein with extremely high affinity *(K_d_* of 0.8-1.6 nM as measured by surface plasmon resonance) (Anderson et al., 2021). The four sdAbs show no obvious similarity in their three variable complementarity-determining regions (CDRs; **Figure 1b-c**), suggesting that they likely each recognize distinct epitopes on the N protein.

**Figure 1.**
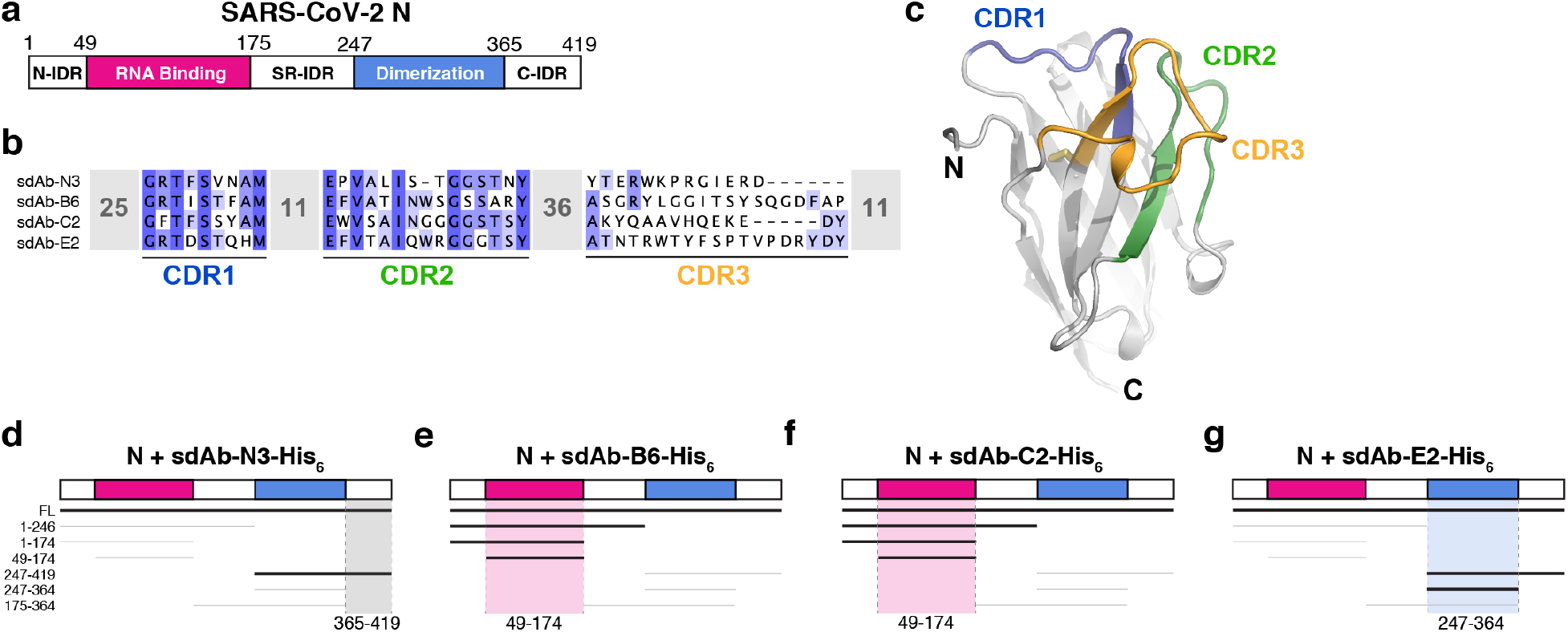
sdAbs target individual domains of the SARS-CoV-2 N protein. (a) Domain schematic of SARS-CoV-2 N, with the RNA-binding domain in magenta, the dimerization domain in blue, and the three intrinsically disordered domains (N-IDR, serine/arginine rich (SR)-ID, and C-IDR) in white. (b) Sequences of four sdAbs targeting the SARS-CoV-2 N protein (Anderson et al., 2021; Sherwood and Hayhurst, 2021) showing the variable sequences within the Complementarity-Determining Regions (CDRs). (c) Example sdAb structure (sdAb-B6 shown) with complementarity-determining regions (CDR) 1, 2, and 3 colored blue, green, and orange, respectively. (d-g) Summary of Ni^2+^ pulldown assays using C-terminally His_6_-tagged sdAbs and seven truncations of SARS-CoV-2 N (see **Figure S1**). Thick lines indicate a positive interaction; highlighted regions indicate inferred region of SARS-CoV-2 N targeted by each sdAb.

The SARS-CoV-2 N protein possesses a modular structure with an N-terminal RNA-binding domain (RBD) and a C-terminal dimerization domain (CTD), plus three intrinsically disordered regions at the N- and C-termini (N-IDR and C-IDR, respectively) and in the serine/arginine region separating the RBD and CTD (SR-IDR; **Figure 1a**). We and others have previously shown that the N protein oligomerizes in vitro through its CTD and disordered C-terminal tail (Ye et al., 2020), and that the protein undergoes liquid-liquid phase separation with RNA mediated by its RBD and central disordered region (Carlson et al., 2020; Cubuk et al., 2020; Iserman et al., 2020; Perdikari et al., 2020; Savastano et al., 2020; Lu et al., 2021; Luo et al., 2021). To determine which domain(s) of N are recognized by each sdAb, we performed direct Ni^2+^ affinity pulldown assays using His_6_-tagged sdAbs and seven N protein truncations spanning its five distinct domains (**Figure 1d-g, Figure S1**). As expected, we found that all four sdAbs robustly bind full-length N protein. We could further narrow the binding region of each sdAb to a defined subdomain: sdAb-B6 and sdAb-C2 each recognize the N protein RBD (residues 49-174), while sdAb-E2 recognizes the CTD (residues 247-364) and sdAb-N3 recognizes the C-IDR (residues 365-419; **Figure 1d-g, Figure S1**).

### Structures of sdAb-N protein complexes

We next sought to determine high-resolution structures of N:sdAb complexes. We first co-crystallized sdAbs C2 and B6 with the N protein RNA-binding domain (residues 49-174; referred to hereafter as N^RBD^), and determined structures of these complexes to 1.42 Å and 2.56 *Å* resolution, respectively (**Table S2**). The structures reveal that the two sdAbs recognize distinct surfaces on the N protein RBD, away from the RNA binding surface. sdAb-C2 recognizes a concave surface on the N protein RBD, burying 765 Å^2^ of surface on each partner (**Figure 2b-c**). The binding interface is made up mostly of the CDR2 (40% of the interface) and CDR3 (53%) loops, with only a minor contribution from CDR1 (7%). The bulk of the interface is hydrophobic, with sdAb-C2 residues A33, W47, A50, Y99, A101, and V103 docking against the N^RBD^ surface. Specific hydrogen bonds are made by sdAb-C2 residues S57 and Q105.

**Figure 2.**
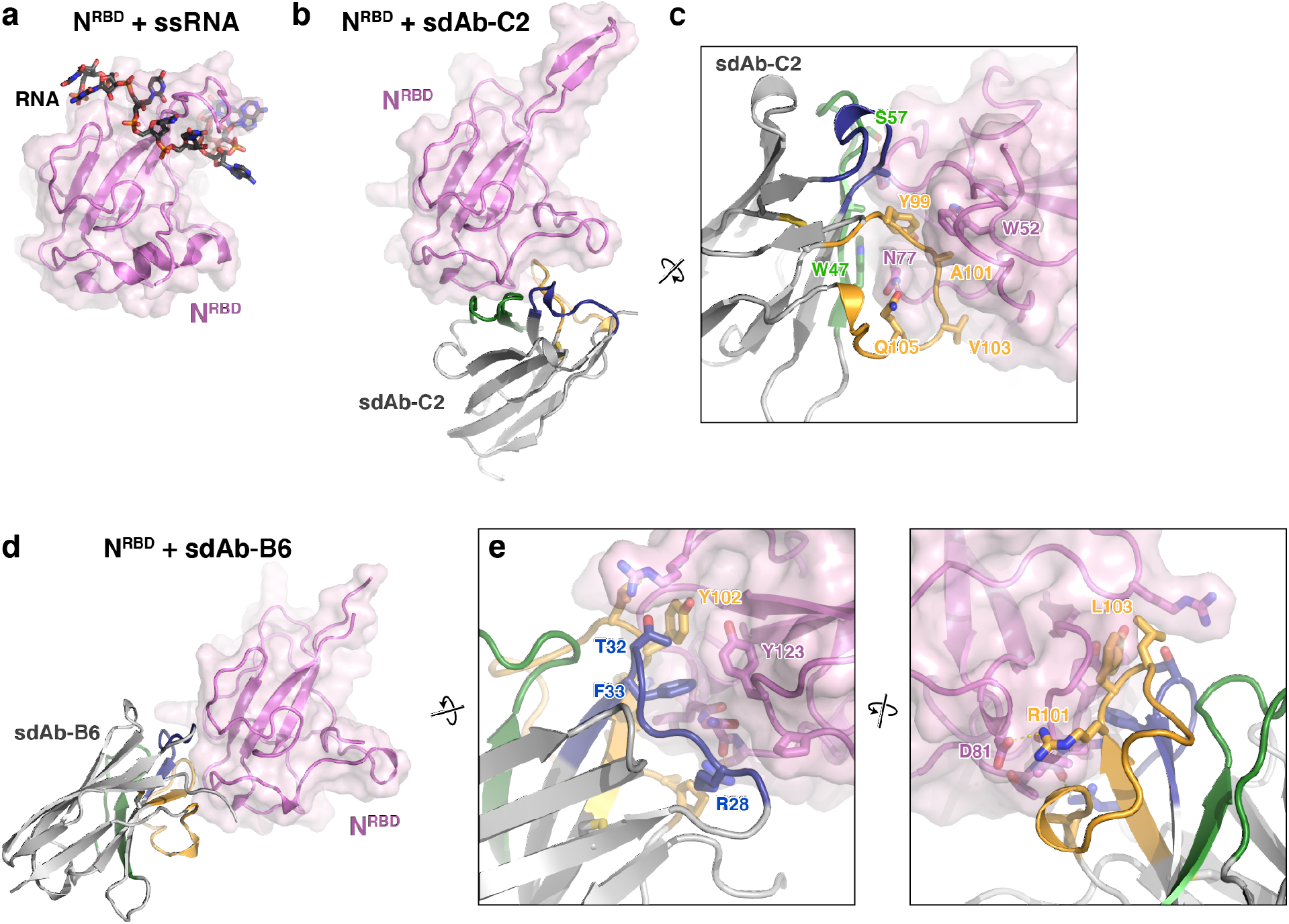
Structures of two sdAbs targeting the SARS-CoV-2 N protein RBD. (a) Structure of the SARS-CoV-2 N^RBD^ (pink) bound to single-stranded RNA (black) (PDB ID 7ACT; (Dinesh et al., 2020)). Panels (b) and (d) show the N^RBD^ in the same orientation. (b) Structure of N^RBD^ (pink) bound to sdAb-C2. CDR1, CDR2, and CDR3 are colored blue, green, and orange as in **Figure 1c**. (c) Closeup view of the N^RBD^:sdAb-C2 complex. (d) Structure of N^RBD^ (pink) bound to sdAb-B6. CDR1, CDR2, and CDR3 are colored blue, green, and orange as in **Figure 1c**. (e) Two closeup views of the N^RBD^:sdAb-B6 complex.

sdAb-B6 binds a separate surface on N^RBD^, burying 617 Å^2^ of surface area on each partner (**Figure 2d-e**). The interface of sdAb-B6 with N is made up mostly of CDR1 (34% of the interface) and CDR3 (60%), with only a minor contribution from CDR2 (4%). At the core of the interface is a pair of aromatic side-chains from sdAb-B6, F33 and Y102, which together dock into a hydrophobic surface on N^RBD^. Two arginine residues, R28 and R101, also make specific hydrogen bonds with N^RBD^.

Next, we co-crystallized sdAb-E2 with the N protein CTD. Initial crystals with the full dimerization domain (residues 247-364) failed to diffract x-rays, so we instead cocrystallized sdAb-E2 with a truncated construct lacking 22 N-terminal residues (residues 269-364; referred to hereafter as N^CTD^). We determined the structure of the N^CTD^:sdAb-E2 complex to 2.2 Å resolution in a crystal form with four independent copies of the N^CTD^ dimer, with each dimer bound to a single sdAb-E2 (**Figure 3a**). All four copies show an identical interface, with sdAb-E2 binding across the flat β-sheet making up the N^CTD^ dimer interface. The interface buries a total of 710 Å^2^ of surface area on each partner, with CDR2 and CDR3 making up nearly the entire interface (33% and 64% of the interface, respectively). CDR1 structurally stabilizes CDR2 and CDR3, but makes no direct contact with the N protein (**Figure 3b**). The majority of the interface involves three aromatic residues on CDR3 – W102, Y104, and F105 – with W102 and Y104 laying nearly flat against the N^CTD^ β-sheet. sdAb-E2 also makes specific hydrogen bonds with Q281 of both N^CTD^ protomers, with one Q281 residue making a salt-bridge interaction with both the main chain and side chain of sdAb-E2 S106, and the other Q281 hydrogen bonding sdAb-E2 residue R54 (**Figure 3b**).

**Figure 3.**
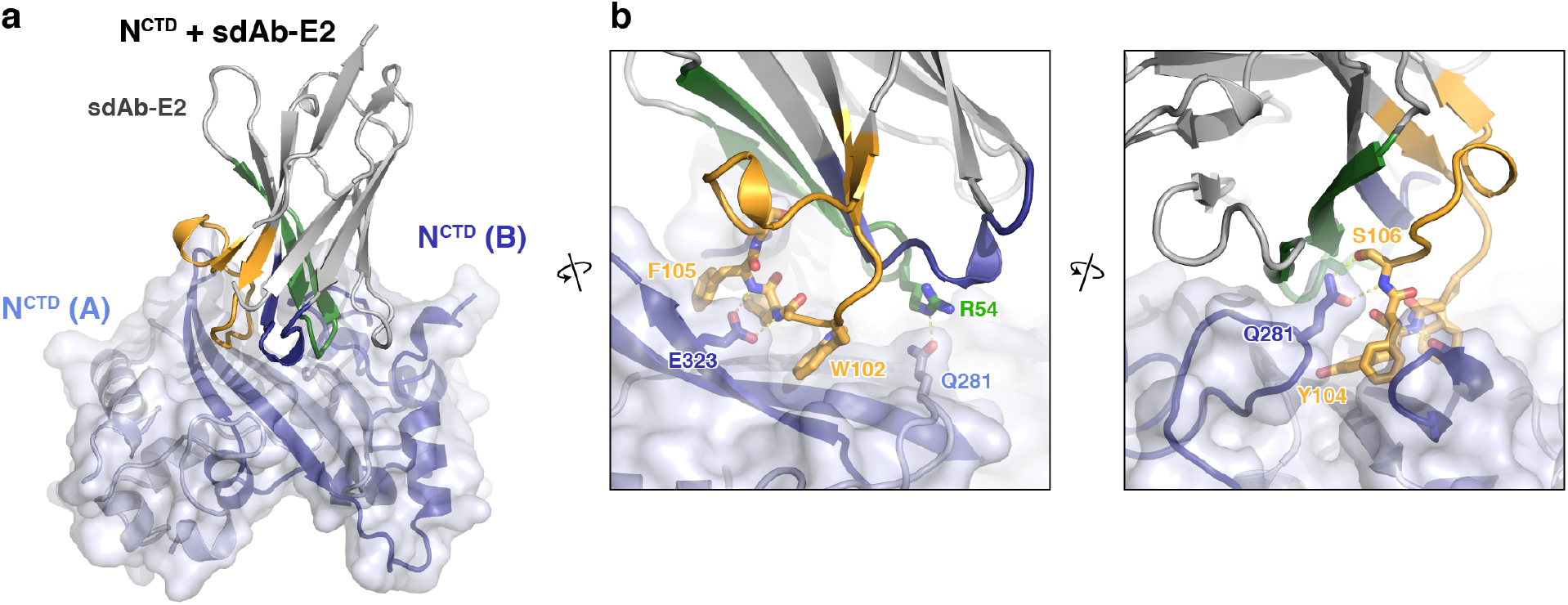
Structure of an sdAb targeting the SARS-CoV-2 N protein CTD. (a) Structure of sdAb-E2 (with CDR1, CDR2, and CDR3 colored blue, green, and orange as in **Figure 1c**) bound to the N^CTD^ dimer (light blue/dark blue). (b) Two closeup views of the N^CTD^:sdAb-E2 complex.

### Variability of sdAb-recognition surfaces on SARS-CoV-2 N

To date, more than 500,000 individual isolates of SARS-CoV-2 have been fully sequenced, enabling a comprehensive examination of variability across the surface of each of its proteins. The sequences of many variants of concern, many of which have higher infectivity than the ancestral SARS-CoV-2 strain, are also known. To determine the likely effectiveness of each sdAb in recognizing variants of concern and the broader population of SARS-CoV-2 variants, we compared the epitopes of each sdAb with a dataset of variants in 489,605 SARS-CoV-2 sequences from the GISAID Initiative (http://www.gisaid.org) and the COVID CG project (http://covidcg.org). These data reveal that the proteins’ three intrinsically disordered regions are mutated more than 20-fold more often than the ordered N^RBD^ or N^CTD^ domains (0.012 mutations/residue/isolate for intrinsically disordered regions vs. 0.0005 mutations/residue/isolate for the ordered domains; **Figure S2a** and **Table S3**), demonstrating strong selective pressure to maintain these domains’ sequence and structure. We also examined N protein mutations in the major SARS-CoV-2 variants of concern worldwide. As with the larger dataset, mutations in these variants strongly cluster in the central intrinsically disordered region, with a few mutations in the N^RBD^ domain (**Figure S2a**).

We compared the epitopes of sdAbs B6, C2, and E2 to the data on N protein variation, and found that the three sdAbs largely bind highly conserved surfaces (**Figure S2b-c**). For sdAb-B6, the most highly-mutated residues in its epitope are N^RBD^ residues T135, L139, and T141, which are mutated in only 1372 (0.28%), 976 (0.19%), and 402 (0.08%) of the 489,605 SARS-CoV-2 genome samples, respectively. The sdAb-C2 epitope overlaps two residues on N^RBD^, D144 and H145, which are mutated in 575 (0.12%) and 2010 (0.41%) SARS-CoV-2 genome samples, respectively. Both sdAb-B6 and sdAb-C2 contact residue P80, which is mutated to arginine in variant P1. This residue is at the edge of both interfaces, however, and its mutation is unlikely to affect N^RBD^ recognition by either sdAb.

On the N^CTD^, sdAb-E2 contacts three residues – T325, S327, and T334 – that are mutated in 373 (0.08%), 511 (0.10%), and 990 (0.20%) SARS-CoV-2 genome samples, respectively. No major variants of concern show mutations in the N^CTD^, illustrating the high conservation of this domain and the potential utility of sdAb-E2 for robust N protein recognition. Overall, this analysis suggests that all three sdAbs will recognize the vast majority of SARS-CoV-2 N proteins in active infections, including those encoded by the major variants of concern worldwide. Given the higher variability of the IDR regions compared to the RBD or CTD, sdAb-N3 (which binds the C-IDR) may be less broadly effective at recognizing N protein variants than the three other sdAbs.

### sdAb binding compromises RNA binding and phase separation of SARS-CoV-2 N

The multiple roles of SARS-CoV-2 N in the viral life cycle rely on its ability to bind RNA and self-assemble through liquid-liquid phase separation. We tested whether addition of sdAbs would affect these biochemical properties, in the context of the full-length N protein. We first tested RNA binding, using electrophoretic mobility shift assays (EMSAs) with short single-stranded or double-stranded RNAs. Structures of the N protein RBD were recently determined by NMR in complex with either single- and double-stranded RNA, revealing distinct binding modes on the same positively-charged surface of the protein (**Figure 2a**) (Dinesh et al., 2020). Using short 7-base single- or double-stranded RNAs also used in this prior study, we performed EMSAs using fulllength N protein in the presence or absence of sdAbs B6 and C2. Despite the use of short RNAs that each can only bind a single N protomer, we found that all binding curves are best fit using a cooperative binding model with Hill coefficient ~5, hinting that binding of even short RNAs is intimately linked with self-assembly of the N protein (**Figure 4a-b**). For full-length N, we measured the binding affinity for ssRNA at 0.56 +/− 0.06 μM, and for dsRNA at 0.48 +/− 0.03 μM, comparable to the affinity measured previously in low-salt buffer (Dinesh et al., 2020). Instead of showing a simple shift from an unbound to a bound band as expected for a 1:1 complex, our gels clearly showed a progressive shift toward the well of the gel at higher N protein concentration, consistent with higher-order assembly of the N-RNA complexes (**Figure S3**). We next added equimolar amounts of either sdAb-B6 or sdAb-C2, both of which bind the N protein RBD. Despite neither sdAb’s epitope overlapping the RNA binding site directly, both sdAbs slightly reduced the affinity for ssRNA (to 0.70 +/− 0.04 and 0.63 +/− 0.04 μM, respectively, for sdAb-B6 and sdAb-C2) (**Figure 4a**). Curiously, while sdAb-C2 had no effect on dsRNA binding (*K_d_*,=0.49 +/− 0.06 μM), sdAb-B6 significantly reduced dsRNA binding (*K_d_*=0.82 +/− 0.11 μM) (**Figure 4b**). Both sdAbs B6 and C2 significantly reduced the progressive band-shift observed with wild-type N protein, especially for the dsRNA binding assays (**Figure S3**). This observation suggests that the two sdAbs affect the N protein-RNA interaction not by interfering with RNA binding of a single N protein protomer, but rather by compromising large-scale self-assembly of N.

**Figure 4.**
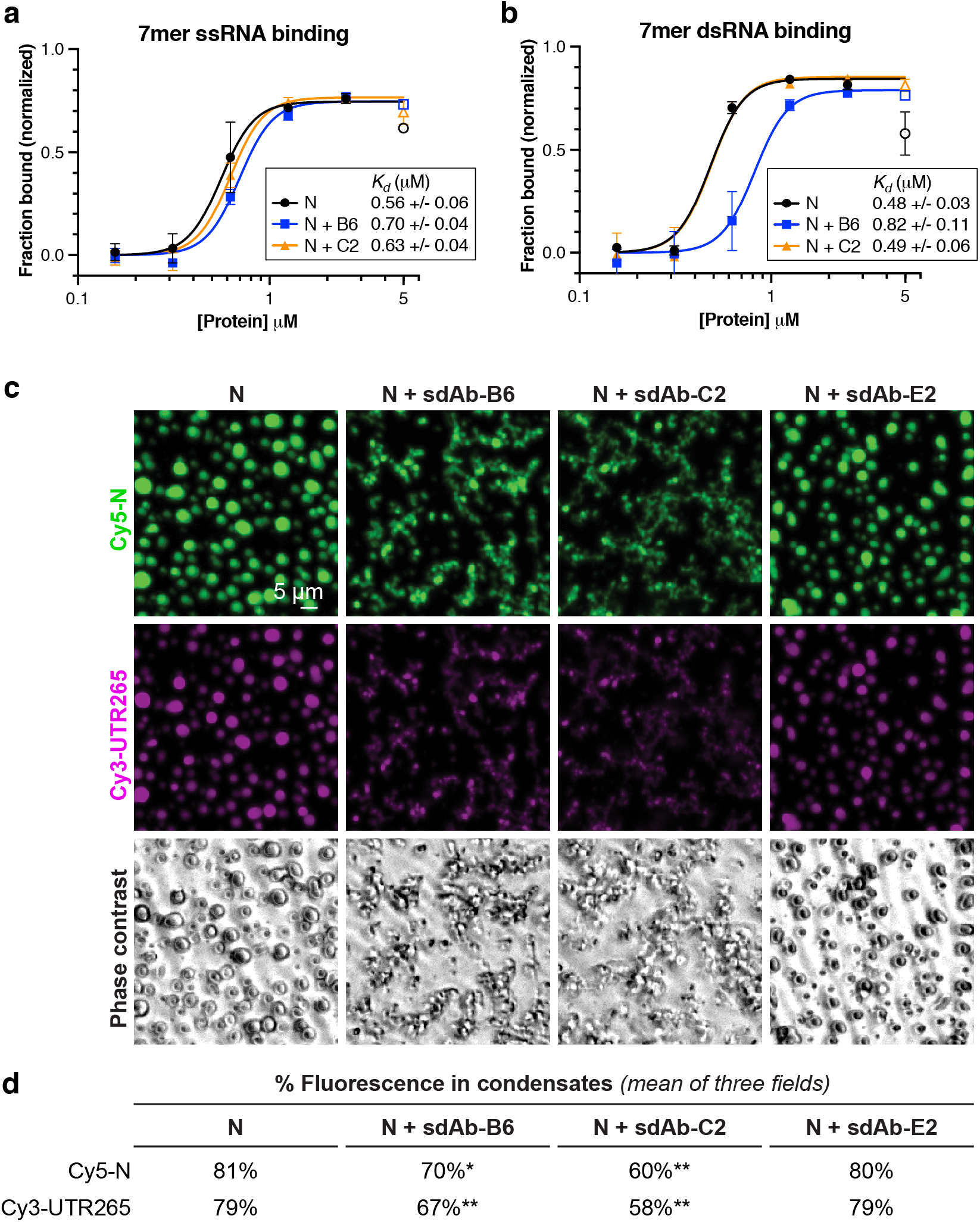
RBD-targeting sdAbs targeting affect RNA binding affinity and RNA-mediated phase separation of SARS-CoV-2 N. (a) Binding curves for SARS-CoV-2 N (full-length protein) binding to a 7-mer singlestranded RNA, in the absence of added sdAbs (black points) or in the presence of equimolar sdAb-B6 (blue points) or sdAb-C2 (orange points). See **Figure S3a** for example electrophoretic mobility shift (EMSA) gels. (b) Binding curves for SARS-CoV-2 N binding to a 7-mer double-stranded RNA, in the absence of added sdAbs (black points) or in the presence of equimolar sdAb-B6 (blue points) or sdAb-C2 (orange points). See **Figure S3c** for example electrophoretic mobility shift (EMSA) gels. (c) In vitro phase separation of Cy5-labeled N (green) and Cy3-labeled UTR265 RNA (magenta), in the absence or presence of equimolar sdAb-B6, sdAb-C2, or sdAb-E2. (d) Quantitation of fluorescence (protein and RNA) in condensates. All measurements represent the mean of three separate microscope fields. Asterisks indicate P-values from unpaired t-tests: *, P < 0.05; **, P < 0.01.

Our RNA binding results suggested that sdAbs may compromise self-assembly of N into large-scale oligomers, including the condensates that we and others have observed both in vitro and in cells. To directly test whether sdAbs compromise LLPS of the N protein, we mixed Cy5-labeled full-length N with a 265-base RNA derived from the 5’ UTR of SARS-CoV-2 (UTR265), and directly observed the formation of phase-separated condensates using light microscopy (**Figure 4c-d**). We observed robust assembly of N+RNA condensates, that was significantly compromised by addition of equimolar amounts of either sdAb-B6 or sdAb-C2 (**Figure 4d**). In addition to reducing the overall amount of LLPS, both sdAbs altered the morphology of the observed condensates, from round droplets to more extended, fibrous structures (**Figure 4c**). Addition of sdAb-E2, which binds the CTD, did not strongly affect formation of N+RNA condensates.

## Discussion

Here, we present high-resolution structures of three single-domain antibodies bound to defined domains of the SARS-CoV-2 N protein. Two sdAbs bind the N-terminal RNA binding domain, and one binds the C-terminal dimerization domain. All three sdAbs recognize highly-conserved surfaces on the N protein, suggesting that they could be used to recognize SARS-CoV-2 infections in antigen tests. As SARS-CoV-2 transitions from an acute public health threat to an endemic phase, the ability to robustly detect and respond to infections and local outbreaks will remain important, especially in the developing world. Development of antigen tests with the sdAbs characterized here, which can be produced at high levels in *E. coli* and recognize the highly-abundant N protein with high specificity and affinity, can significantly aid this effort.

## Materials and Methods

### Cloning and protein purification

We previously reported the cloning, expression, and purification of full-length and truncated SARS-CoV-2 N protein (Ye et al., 2020; Lu et al., 2021). Briefly, constructs were amplified by PCR from the IDT 2019-nCoV N positive control plasmid (IDT cat. # 10006625; NCBI RefSeq YP_009724397) and inserted by ligation-independent cloning into UC Berkeley Macrolab vectors 2B-T (AmpR, N-terminal His_6_-fusion; Addgene #29666) or 2C-T (AmpR, N-terminal His_6_-MBP fusion; Addgene #29706). Plasmids were transformed into *E. coli* strain Rosetta 2(DE3) pLysS (Novagen), and grown in the presence of ampicillin and chloramphenicol to an OD_600_ of 0.8 at 37°C, induced with 0.25 mM IPTG, then grown for a further 16 hours at 18°C prior to harvesting by centrifugation. Harvested cells were resuspended in buffer A (25 mM Tris-HCl pH 7.5, 5 mM MgCl_2_ 10% glycerol, 5 mM β-mercaptoethanol, 1 mM NaN_3_) plus 500 mM NaCl (for full-length N, 1 M NaCl) and 5 mM imidazole pH 8.0. For purification, cells were lysed by sonication, then clarified lysates were loaded onto a Ni^2+^ affinity column (Ni-NTA Superflow; Qiagen), washed in buffer A plus 300 mM NaCl and 20 mM imidazole pH 8.0, and eluted in buffer A plus 300 mM NaCl and 400 mM imidazole. For cleavage of His_6_-tags, proteins were buffer exchanged in centrifugal concentrators (Amicon Ultra, EMD Millipore) to buffer A plus 300 mM NaCl and 20 mM imidazole, then incubated 16 hours at 4°C with TEV protease (Tropea et al., 2009). Cleavage reactions were passed through a Ni^2+^ affinity column again to remove uncleaved protein, cleaved His_6_-tags, and His_6_-tagged TEV protease. Proteins were concentrated in centrifugal concentrators and purified by size-exclusion chromatography (Superdex 200; GE Life Sciences) in gel filtration buffer (25 mM Tris-HCl pH 7.5, 300 mM NaCl, 5 mM MgCl_2_, 10% glycerol, 1 mM DTT). Purified proteins were concentrated and stored at 4°C for analysis.

Vectors encoding sdAbs B6, C2, and E2 in pET22b were generously provided by Ellen Goldman (Anderson et al., 2021). These vectors encode each sdAb with a pelB leader sequence at the N-terminus for periplasmic expression, and a His_6_-tag at the C-terminus for purification (**Table S1**). For sdAb-N3, a codon-optimized gene was synthesized (Integrated DNA Technologies) and inserted into pET22b. All sdAbs were transformed into Rosetta 2(DE3) pLysS, grown in the presence of ampicillin and chloramphenicol to an OD_600_ of 0.8 at 37°C, induced with 0.5 mM IPTG, then grown for a further 4 hours at 24°C prior to harvesting by centrifugation. Proteins were purified as above using Ni^2+^ affinity and size exclusion chromatography (Superdex 75, GE Life Sciences), excluding cleavage of the His_6_-tags.

### Nickel pulldown assays

For Ni^2+^ affinity pulldowns, 10 μg sdAb-His_6_ fusion protein was mixed with 10 μg N protein (FL or truncated) in a total reaction volume of 50 μl, in binding buffer (20 mM HEPES pH 7.5, 200 mM NaCl, 5 mM MgCl_2_, 5% glycerol, 10 mM Imidazole, and 0.1% NP-40). After a 60 minute incubation at room temperature, 30 μl Ni^2+^ beads equilibrated in binding buffer (Qiagen Ni-NTA superflow) was added, and the mixture incubated a further 20 minutes at room temperature with rotation. Beads were then washed 3X with 1 mL binding buffer, and bound proteins were eluted by adding 5 μL of 2 M Imidazole plus 20 μL of 2X Laemmli sample buffer. Input (5 μl of the initial reaction) and bound (15 μl of elution fraction) samples were analyzed by SDS-PAGE and Coomassie blue staining.

### Crystallography

For crystallization of the N^49-174^:sdAb-C2 complex, sdAb-C2 and N^49-174^ were exchanged into crystallization buffer (20 mM HEPES pH 7.0, 200 mM NaCl, 5 mM MgCl_2_, 1 mM TCEP) and mixed at a 1:1 molar ratio of sdAb to N, for a final protein concentration of 15 mg/mL. Crystals were obtained in sitting drop format by mixing protein 1:1 with well solution containing 0.1 M Tris-HCl pH 8.5, 0.2 M LiSO4, and 20% PEG 4000. Crystals were transferred to a cryoprotectant containing an additional 10% glycerol, then flash-frozen in liquid nitrogen. Diffraction data were collected at Advanced Photon Source, NE-CAT beamline 24ID-E (support statement below). Data was automatically indexed and reduced by the RAPD data-processing pipeline (https://github.com/RAPD/RAPD), which uses XDS (Kabsch, 2010) for indexing and integration, and the CCP4 programs AIMLESS and TRUNCATE (Winn et al., 2011) for scaling and structure-factor calculation. The structure was determined by molecular replacement using PHASER (McCoy et al., 2007) with a structure of the SARS-CoV-2 RNA binding domain (PDB ID 7CDZ; (Peng et al., 2020)) and a structure of an alpaca nanobody selected against the Hepatitis C virus E2 glycoprotein (PDB ID 4JVP; (Tarr et al., 2013)). Models were manually rebuilt in COOT (Emsley et al., 2010) and refined in phenix.refine (Afonine et al., 2012) (**Table S2**).

For crystallization of the N^49-174^:sdAb-B6 complex, sdAb-B6 and N^49-174^ were exchanged into crystallization buffer (20 mM HEPES pH 7.5, 200 mM NaCl, 5 mM MgCl_2_, 1 mM TCEP) and mixed at a 1:1 molar ratio of sdAb to N, for a final protein concentration of 15 mg/mL. Crystals were obtained in hanging drop format by mixing protein 1:1 with well solution containing 0.1 M sodium citrate pH 5.6, 0.1 M sodium-potassium tartrate, and 19% PEG 3350. Crystals were transferred to a cryoprotectant containing an additional 10% glycerol, then flash-frozen in liquid nitrogen. Diffraction data were collected at Advanced Photon Source, NE-CAT beamline 24ID-C (support statement below), and data were processed and the structure determined as above. The structure was determined by molecular replacement using PHASER with separate N^49-174^ and nanobody scaffold chains as above. The model, comprising three identical copies of a 1:1 N^49-174^:sdAb-C2 complex, was manually rebuilt in COOT and refined with phenix.refine (**Table S2**).

For crystallization of the N^269-364^:sdAb-E2 complex, sdAb-E2 and N^269-364^ were separately subjected to surface lysine demethylation by treatment of the purified protein with borane (50 mM final concentration) and formaldehyde (100 mM) for two hours at 4°C, followed by addition of glycine (25 mM final concentration) to quench the reaction, then buffer-exchange by centrifugal concentration and dilution. Methylated N^269-364^ and sdAb-E2 were mixed at a ratio of two N^269-364^ (one dimer) to one sdAb-E2 in crystallization buffer (20 mM HEPES pH 7.0, 200 mM NaCl, 5 mM MgCl_2_, 1 mM TCEP), according to the inferred ratio of these proteins in copurified samples. Crystals were obtained in hanging drop format by mixing protein 1:1 with well solution containing 0.1 M imidazole pH 8.5, 0.1 M calcium acetate, 12% isopropanol, and 24% PEG 3350. Crystals were transferred to a cryoprotectant containing 10% glycerol, then flash-frozen in liquid nitrogen. Diffraction data were collected at Advanced Light Source, Berkeley Center for Structural Biology beamline 5.0.2 (support statement below), and data were processed by XDS (Kabsch, 2010) and the CCP4 programs aimless and truncate (Winn et al., 2011; Evans and Murshudov, 2013). The structure was determined by molecular replacement using PHASER with separate N^269-364^ dimer and nanobody scaffold chains as above. The model, comprising four identical copies of a complex with a dimers N^269-364^ bound to a single sdAb-E2, was manually rebuilt in COOT and refined in phenix.refine. Because of strong diffraction anisotropy and translational non-crystallographic symmetry (tNCS), refinement was performed against a pruned dataset with the lowest-intensity ~17% of reflections removed from the dataset by PHASER in NCS mode, and refined only to 2.2 Å resolution as higher resolution bins showed less than 75% completeness after pruning. Since some protomers of N^269-364^ showed poor electron density in regions distant from the sdAb-E2 interface, we also applied NCS restraints against equivalent chains (A/C/E/G (N^269-364^), B/D/F/H (N^269-364^), and I/J/K/L(sdAb-E2)) to maintain stability in the refinement (**Table S2**).

### Sequence and structure analysis

For examination of N protein variation, a dataset of N protein mutations in 489,605 SARS-CoV-2 genomes was downloaded from the COVID CG project (http://covidcg.org) in February 2021. Highly-mutated positions were graphed in Prism v. 9 (Graphpad Software). Mutations in variants of concern were obtained from https://outbreak.info/. Analysis of sdAb-N interfaces was performed with PDBePISA (https://www.ebi.ac.uk/pdbe/pisa/), and graphed alongside N protein variation data.

### RNA binding assays

RNA binding was performed with 5’-fluorescein (FAM) labeled 7mer single-stranded RNA (5’-CACUGAC-3’) or dsRNA (5’-FAM-CACUGAC-3’ annealed with 5’-GUCAGUG-3’ by slow cooling) in a reaction buffer with 20 mM Tris-HCl pH 7.5, 50 mM KCl, 5 mM MgCl_2_, 1 mM DTT, and 5% glycerol. RNA at 100 nM was mixed with protein (5, 2.5, 1.25, 0.625, 0.3125, 0.15625, and 0 μM) in a total reaction buffer of 20 μL and incubated 1 hour at 4°C. 10 μL of each reaction was mixed with 2 μL 6X loading dye, and samples were loaded onto a 10% TBE-PAGE gel (pre-run in 0.5X TBE at 200 volts for 1 hour at 4°C), and run at 150 volts for 40 minutes at 4°C. Gels were imaged on a Bio-Rad ChemiDoc XRS+ imager set to detect Cy-2 with a 0.5 second exposure. Bound/unbound fractions were calculated in ImageJ (Schneider et al., 2012) and binding curves were calculated in Prism v. 9 (Graphpad Software) using a cooperative model with Hill coefficient fixed at 5 to stabilize curve-fitting.

### Phase separation assays

In vitro phase separation assays were performed at 25°C. Unlabeled N protein was mixed with Cy5-labeled protein (linked to an engineered N-terminal cysteine using maleimide linkage) at 1:10 ratio, then mixed with each sdAb at 1:1 ratio. Phase separation of 40 μM N protein (in 5 mM HEPES pH 7.5, 160 mM KCl) was induced by adding to a pre-mixture of three times volumes of UTR265 RNA (13.3 ng/μL in 5 mM HEPES pH 7.5). Samples were mixed in protein LoBind tubes (Eppendorf, 022431064) and then immediately transferred onto 18-well glass-bottom chamber slides (iBidi, 81817). Condensates were imaged after 40 min when the phase separation particles are stable. Images were taken on a CQ1 confocal quantitative image microscope (Yokogawa) with 20x-PH objective. For quantitation, condensates were identified by particle analysis in CQ1 analysis software, then the total fluorescence inside condensates and the total volume of condensates were calculated and divided by the total fluorescence in the field or the total volume of the field to yield a percentage value.

>UTR265 (nt 1-265 of SARS-CoV-2 genome) AUUAAAGGUUUAUACCUUCCCAGGUAACAAACCAACCAACUUUCGAUCUCUUGUAGAUCUGUUCUCUAAACGAACUUUAAAAUCUGUGUGGCUGUCACUCGGCUGCAUGCUUAGUGCACUCACGCAGUAUAAUUAAUAACUAAUUACUGUCGUUGACAGGACACGAGUAACUCGUCUAUCUUCUGCAGGCUGCUUACGGUUUCGUCCGUGUUGCAGCCGAUCAUCAGCACAUCUAGGUUUCGUCCGGGUGUGACCGAAAGGUAAG

### Beamline support statements

This work is based upon research conducted at the Northeastern Collaborative Access Team beamlines at the Advanced Photon Source, which are funded by the National Institute of General Medical Sciences from the National Institutes of Health (P30 GM124165). The Eiger 16M detector on 24-ID-E is funded by a NIH-ORIP HEI grant (S10OD021527). This research used resources of the Advanced Photon Source, a U.S. Department of Energy (DOE) Office of Science User Facility operated for the DOE Office of Science by Argonne National Laboratory under Contract No. DE-AC02-06CH11357. The Berkeley Center for Structural Biology is supported in part by the Howard Hughes Medical Institute. The Advanced Light Source is a Department of Energy Office of Science User Facility under Contract No. DE-AC02-05CH11231. The ALS-ENABLE beamlines are supported in part by the National Institutes of Health, National Institute of General Medical Sciences, grant P30 GM124169.

## Acknowledgements

The authors thank Ellen Goldman for the kind gift of vectors encoding sdAb-B6, sdAb-C2, and sdAb-E2; Stacey Ortega and Marc Allaire at Advanced Light Source beamline 5.0.2; the staff of Advanced Light Source NE-CAT beamlines 24ID-C and 24ID-E; Haiyang Yu for helpful experimental suggestions, and members of the Corbett lab for critical feedback. KDC acknowledges support from UC San Diego and the National Institutes of Health (R01GM128464).

