## Supplementary material for "Structural basis for SARS-CoV-2 Nucleocapsid protein recognition by single-domain antibodies": Tables S1-S2, Figures S1-S3

### Table S1: Protein sequences

| Protein | GenBank ID | Sequence |
|---|---|---|
| SARS-CoV-2 N | YP_009724397 | MSDNGPQNQRNAPRITFGGPSDSTGSNQNGERSGARSKQRRPQGLPNNTASW<br>FTALTQHGKEDLKFPRGQGVPINTNSSPDDQIGYYRRATRRIRGGDGKMKDL<br>SPRWYFYYLGTGPEAGLPYGANKDGIIWVATEGALNTPKDHIGTRNPANNAA<br>IVLQLPQGTTLPKGFYAEGSRGGSQASSRSSSRSRNSSRNSTPGSSRGTSPA<br>RMAGNGGDAALALLLLDRLNQLESKMSGKGQQQQGQTVTKKSAAEASKKPRQ<br>KRTATKAYNVTQAFGRRGPEQTQGNFGDQELIRQGTDYKHWPQIAQFAPSAS<br>AFFGMSRIGMEVTPSGTWLTYTGAIKLDDKDPNFKDQVILLNKHIDAYKTFP<br>PTEPKKDKKKKADETQALPQRQKKQQTVTLLPAADLDDFSKQLQQSMSSADS<br>TQA |
| sdAb-B6 | N/A | MAEVQLQASGGGLVQAGDSLRLSCVAVSGRTISTFAMGWFRQAPGKEREFVA<br>TINWSGSSARYADPVEGRFTISRDDAKNTVYLEMSSLKPGDSAVYYCASGRY<br>LGGITSYSQGDFAPWGQGTQVTVSSAAALEHHHHHH |
| sdAb-C2 | N/A | MAEVQLQASGGGLVRPGGSLRLSCAASGFTFSSYAMMWVRQAPGKGLEWVSA<br>INGGGGSTSYADSVKGRFTISRDNAKNTLYLQMNSLKPEDTAVYYCAKYQAA<br>VHQEKEDYWGQGTQVTVSSAAALEHHHHHH |
| sdAb-E2 | N/A | MAEVQLQASGGGLVQAGGSLRLSCAASGRTDSTQHMAWFRQAPGKEREFVTA<br>IQWRGGGTSYTDSVKGRFTISRDNAKNTVYLEMNSLKPEDTAVYYCATNTRW<br>TYFSPTVPDRYDYWGQGTQVTVSSAAALEHHHHHH |
| sdAb-N3 | N/A | MAKVKLQQSGGGLVQTGGSLRLSCAPSGRTFSVNAMGWYRQAPGKQREPVAL<br>ISTGGSTNYADSVKGRFTISRDNVKNTVYLQMNTLKPEDTAVYYCYTERWKP<br>RGIERDWGQGTQVTVSSAAALEHHHHH |

[1]Red text shows the N-terminal MA sequence remaining after signal peptide cleavage, and the C-terminal $His_6$-tag used for purification of sdAbs.

### Table S2: Data collection and refinement statistics

| Data collection | sdAb-B6:N$^{49-174}$ | sdAb-C2:N$^{49-174}$ | sdAb-E2:N$^{269-364}$ |
|---|---|---|---|
| Beamline | APS 24ID-C | APS 24ID-E | ALS 5.0.2 |
| Data collection date | March 24, 2021 | February 11, 2021 | May 14, 2021 |
| Resolution (Å) | 112 – 2.56 | 77 – 1.42 | 50 – 1.80 |
| Wavelength (Å) | 0.97918 | 0.97918 | 1.00004 |
| Space Group | $C222_1$ | $P2_1$ | $P2_12_12_1$ |
| Unit Cell Dimensions (a, b, c) Å | 99.87, 172.98, 112.37 | 46.67, 77.12, 71.93 | 75.85, 131.556, 140.05 |
| Unit cell Angles ($\alpha,\beta,\gamma$) ° | 90, 90, 90 | 90, 95.14, 90 | 90, 90, 90 |
| $I/\sigma$ (last shell) | 3.2 (0.4) | 25.7 (4.5) | 8.1 (0.4) |
| [1]$R_{sym}$ (last shell) | 0.259 (3.807) | 0.037 (0.281) | 0.089 (5.417) |
| [2]$R_{meas}$ (last shell) | 0.269 (3.947) | 0.041 (0.307) | 0.096 (5.884) |
| [3]CC1/2 (last shell) | | | |
| Completeness (last shell) % | 99.6 (99.9) | 99.1 (94.4) | 99.7 (99.5) |
| Number of reflections | 430193 | 632696 | 850156 |
| *unique* | 31608 | 95122 | 129860 |
| Multiplicity (last shell) | 13.6 (14.3) | 6.7 (6.2) | 6.5 (6.5) |
| **Refinement** | | | |
| Resolution (Å) | 86.5 – 2.56 | 52.5 – 1.42 | 50 – 2.20 |
| No. of reflections | 29684 | 95054 | 64008 |
| *working* | 28282 | 90426 | 60713 |
| *free* | 1402 | 4628 | 3295 |
| [4]$R_{work}$ (last shell) (%) | 23.36 (41.18) | 14.05 (16.89) | 23.94 (34.11) |
| [4]$R_{free}$ (last shell) (%) | 26.61 (43.43) | 16.62 (19.81) | 27.14 (39.04) |
| **Structure/Stereochemistry** | | | |
| No. of atoms | 10988 | 8295 | 11100 |
| *hydrogen* | 5342 | 3738 | 0 |
| *solvent* | 42 | 672 | 862 |
| *ligand* | 0 | 15 | 5 |
| r.m.s.d. bond lengths (Å) | 0.005 | 0.008 | 0.003 |
| r.m.s.d. bond angles (°) | 0.705 | 0.964 | 0.683 |
| Ramachandran favored/allowed (%) | 98.62/100 | 98.37/100 | 99.61/100 |
| MolProbity Clashscore/Overall score | 7.67/1.37 | 2.49/1.03 | 4.30/1.40 |
| [5]Protein Data Bank ID | 7N0S | 7N0R | 7N0I |
| [6]SBGrid Data Bank ID | 837 | 836 | 835 |

[1]$R_{sym} = \sum\sum_j |I_j - \langle I \rangle| / \sum I_j$, where $I_j$ is the intensity measurement for reflection j and $\langle I \rangle$ is the mean intensity for multiply recorded reflections.

[2]$R_{meas} = \sum_h [\sqrt{(n/(n-1))} \sum_j [I_{hj} - \langle I_h \rangle] / \sum_{hj} \langle I_h \rangle$

where $I_{hj}$ is a single intensity measurement for reflection h, $\langle I_h \rangle$ is the average intensity measurement for multiply recorded reflections, and $n$ is the number of observations of reflection h.

[3]$CC_{1/2}$ is the Pearson correlation coefficient between the average measured intensities of two randomly-assigned half-sets of the measurements of each unique reflection (Karplus and Diederichs, 2012). $CC_{1/2}$ is considered significant above a value of ~0.15.

[4]$R_{work, free} = \sum||F_{obs}| - |F_{calc}||/|F_{obs}|$, where the working and free $R$-factors are calculated using the working and free reflection sets, respectively.

[5]Coordinates and structure factors have been deposited with the Protein Data Bank (http://www.pdb.org) with the noted accession codes.
[6]Diffraction data have been deposited with the SBGrid Data Bank (http://data.sbgrid.org) with the noted accession codes.

### Table S3: N protein mutations in 489,605 SARS-CoV-2 genomic samples

*See attached Excel spreadsheet*

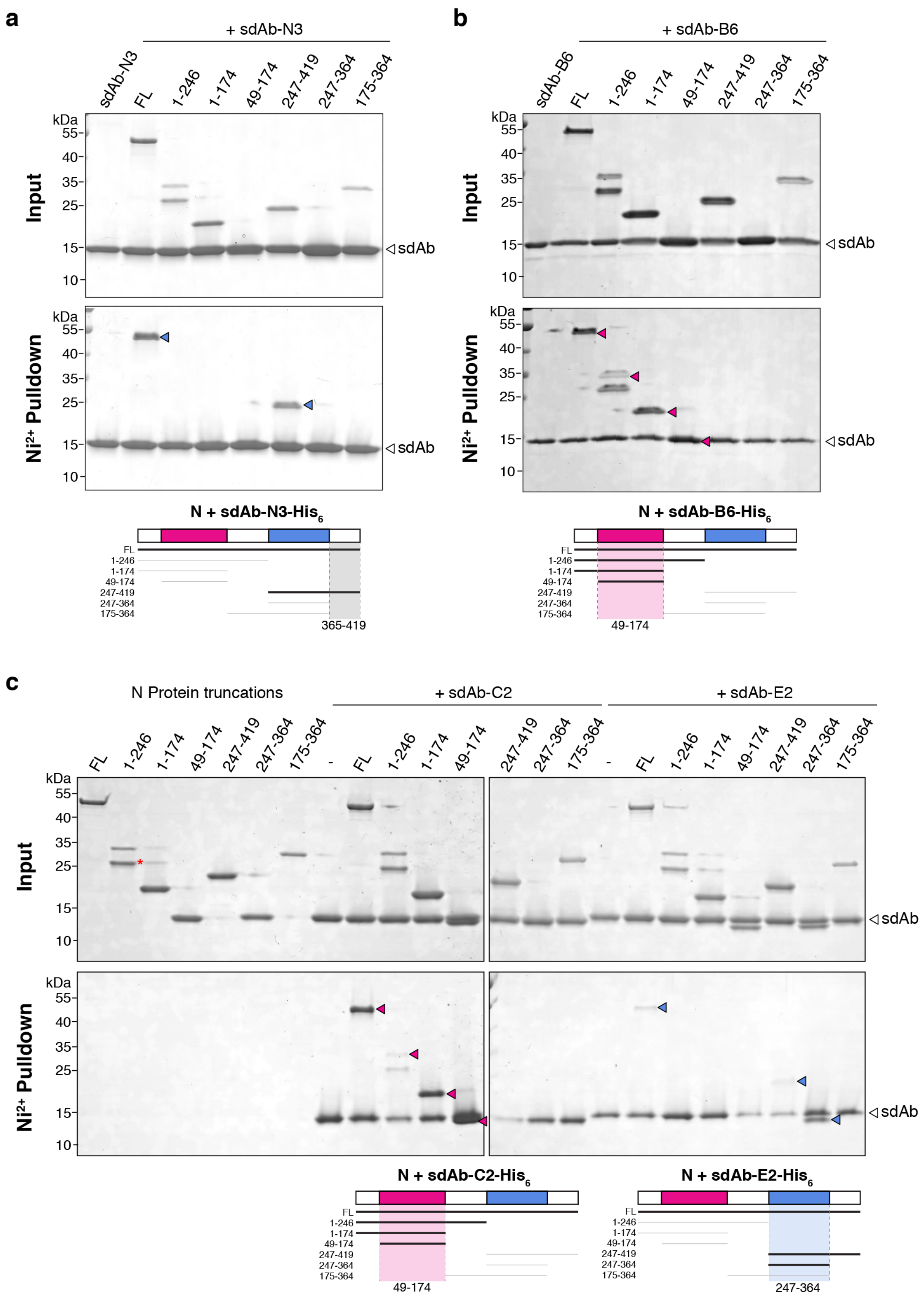
a
+ sdAb-N3
sdAb-N3
FL
1-246
1-174
49-174
247-419
247-364
175-364
kDa
Input
Ni²⁺ Pulldown
sdAb
N + sdAb-N3-His6
365-419
b
+ sdAb-B6
sdAb-B6
N + sdAb-B6-His6
49-174
c
N Protein truncations
+ sdAb-C2
+ sdAb-E2
N + sdAb-C2-His6
N + sdAb-E2-His6
247-364

**Figure S1. sdAbs target individual domains of SARS-CoV-2 N.**

(a) $Ni^{2+}$ pulldown with C-terminally $His_6$-tagged sdAb-N3 and seven truncations of SARS-CoV-2. Positive interactions are shown with blue arrowheads. Schematic (bottom) indicates that sdAb-N3 binds the C-terminal IDR of N (residues 365-419).

(b) $Ni^{2+}$ pulldown with C-terminally $His_6$-tagged sdAb-B6 and seven truncations of SARS-CoV-2. Positive interactions are shown with pink arrowheads. Schematic (bottom) indicates that sdAb-B6 binds the RNA-binding domain of N (residues 49-174).

(c) $Ni^{2+}$ pulldown with C-terminally $His_6$-tagged sdAb-C2 or sdAb-E2 and seven truncations of SARS-CoV-2 (control pulldowns without sdAbs shown at left). Positive interactions are shown with pink (C2) or blue (E2) arrowheads. Schematic (bottom) indicates that sdAb-C2 binds the RNA-binding domain of N (residues 49-174), and sdAb-E2 binds the C-terminal dimerization domain of N (residues 247-364).

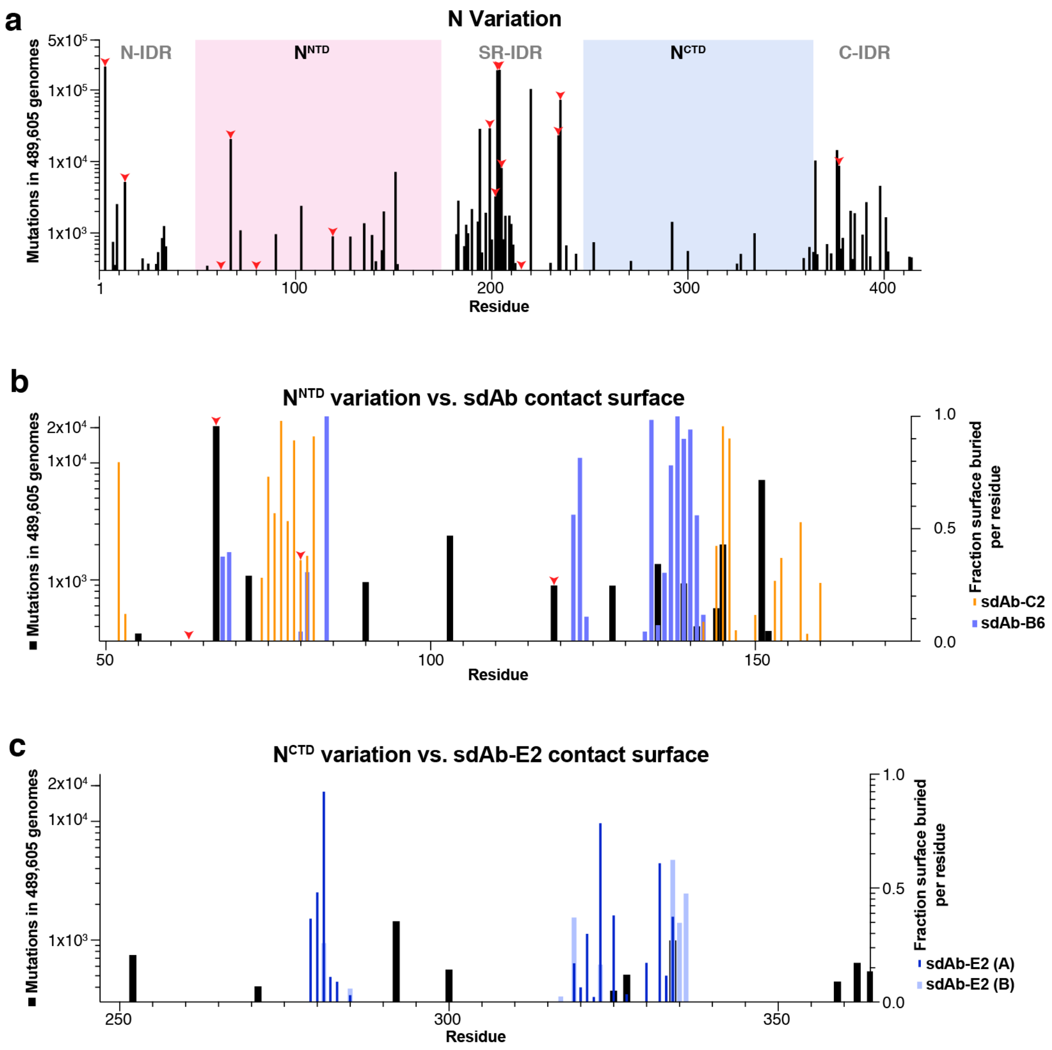


**Figure S2. N protein variation and recognition by sdAbs.**

(a) Histogram showing mutation frequency of N protein in a dataset of 489,605 SARS-CoV-2 genomes (see **Materials and Methods**). N protein domains are labeled, with $N^{RBD}$ highlighted in pink and $N^{CTD}$ highlighted in blue. Red arrowheads indicate positions mutated in one or more variants of concern: B.1.1.7 (D3L, R203K, G204R, S235F), B.1.351 (T205I), B.1.427 (T205I), B.1.429 (T205I), B.1.526 (P199L, M234I), B.1.526.1 (T205I, M234I), B.1.526.2 (P13L, S202R), B.1.617 (del204, del215), B.1.617.1 (R203M, D377Y), B.1.617.2 (D63G, R203M, D377Y), B.1.617.3 (P67S, R203M, D377Y), P.1 (P80R, R203K, G204R), and P.2 (A119S, R203K, G204R, M234I).

(b) N protein variation (black bars, left y axis) versus residue-by-residue buried surface area for the $N^{RBD}$:sdAb-C2 (orange bars, right y axis) and $N^{RBD}$:sdAb-B6 (blue bars, right y axis) complexes.

(c) N protein variation (black bars, left y axis) versus residue-by-residue buried surface area for the $N^{CTD}$:sdAb-E2 complex (blue bars, right y axis). $N^{CTD}$ chain A is shown in dark blue, and chain B in light blue.

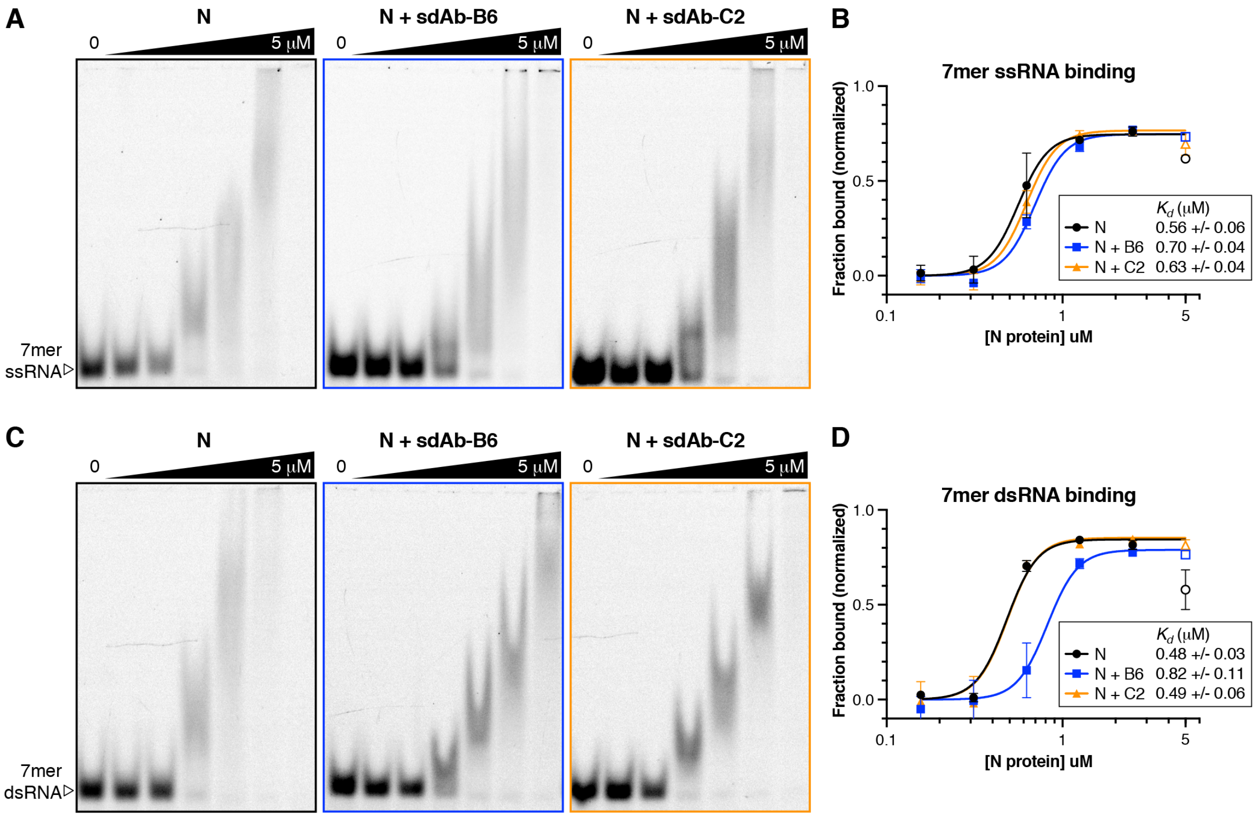


**Figure S3 – Single-domain antibodies affect RNA binding by SARS-CoV-2 N**

(a) Electrophoretic mobility shift assays (EMSA) showing binding of 7mer single-stranded RNA by full-length N protein alone (black boxes) and mixed 1:1 with sdAb-B6 (blue) or sdAb-C2 (orange). Protein concentrations (left to right): 0, 0.15625, 0.3125, 0.625, 1.25, 2.5, and 5 μM.

(b) Binding curves for N protein binding 7mer single-stranded RNA. All measurements were performed in triplicate, and binding curves were fit using a cooperative binding model with a Hill coefficient of 5 (see **Materials and Methods**). Data points at 5 μM protein concentration (outline symbols) were not used for curve fitting because of high error.

(c) EMSA showing binding of 7mer double-stranded RNA by full-length N protein alone (black boxes) and mixed 1:1 with sdAb-B6 (blue) or sdAb-C2 (orange).

(d) Binding curves for N protein binding 7mer double-stranded RNA.
